# Ultrastructural readout of *in vivo* synaptic activity for functional connectomics

**DOI:** 10.1101/2021.07.07.451278

**Authors:** Anna Simon, Arnd Roth, Arlo Sheridan, Mehmet Fişek, Vincenzo Marra, Claudia Racca, Jan Funke, Kevin Staras, Michael Häusser

**Affiliations:** Wolfson Institute for Biomedical Research, University College London, London, UK; HHMI Janelia Research Campus, Ashburn, VA, USA; Salk Institute for Biological Studies, La Jolla, CA, USA; Sussex Neuroscience, University of Sussex, Brighton, UK; University of Leicester, Leicester, UK; Biosciences Institute, Newcastle University, Newcastle upon Tyne, UK

**Keywords:** connectomics, *in vivo*, synapse, electron microscopy, cortex

## Abstract

Large-volume ultrastructural mapping approaches yield detailed circuit wiring diagrams but lack an integrated synaptic activity readout which is essential for functional interpretation of the connectome. Here we resolve this limitation by combining functional synaptic labelling *in vivo* with focused ion-beam scanning electron microscopy (FIBSEM) and machine learning-based segmentation. Our approach generates high-resolution near-isotropic three-dimensional readouts of activated vesicle pools across large populations of individual synapses in a volume of tissue, opening the way for detailed functional connectomics studies. We apply this method to measure presynaptic activity in an ultrastructural context in synapses activated by sensory input in primary visual cortex in awake head-fixed mice, showing that the numbers of recycling and non-recycling vesicles approximate to a lognormal distribution across a large number of synapses. We also demonstrate that neighbouring boutons of the same axon, which share the same spiking activity, can differ greatly in their presynaptic release probability.

## Main

High-throughput, large-volume ultrastructural imaging platforms offer high-resolution data on structural connectivity relationships in brain circuits (Denk and Horstmann, 2004; Kasthuri et al., 2015; Xu et al., 2017). Nonetheless, the lack of parallel functional information on synaptic weights and synaptic activity patterns limits a full understanding of the operation of such circuits (Swanson and Lichtman, 2016). A potential solution to this challenge is to exploit a marker of synaptic activity that can be directly visualized in ultrastructure, allowing functional information to be overlaid on the wiring diagram. FM1-43FX, an established fluorescent reporter of recycled vesicles (Betz and Bewick, 1992; Ryan et al., 1993), meets this demand thanks to its ability to readily photoconvert diaminobenzidine (DAB) to an osmiophilic polymer (Harata et al., 2001; Marra et al., 2012; Rizzoli and Betz, 2004; Schikorski and Stevens, 2001). In cultured neurons and acute brain slice preparations, ultrastructural characterization of this electron-dense vesicular signal has been used to reveal fundamental principles of synaptic size-strength relationships (Branco et al., 2008; Harata et al., 2001; de Lange et al., 2003; Marra et al., 2012; Rey et al., 2020, 2015; Schikorski and Stevens, 2001).

Applying this method to interrogate functional properties across large-scale circuits from intact CNS is more challenging, however. Approaches based on conventional serial section transmission electron microscopy are typically limited to very small tissue volumes and therefore few synaptic terminals. Moreover, thick sections (> 50 nm) limit the accuracy of vesicle detection and classification, generate potential errors of vesicle sampling (Hedreen, 1998) and negate a quasi-continuous readout of vesicle pool organization through the tissue block.

Here we outline an approach that combines *in vivo* FM1-43FX dye-labelling in awake mice, dye-photoconversion, and imaging with FIBSEM, a serial block-face platform that couples a focused ion beam for sample milling with a scanning electron microscope to repetitively image the freshly-exposed block-face, yielding a high-resolution aligned image stack of the tissue volume (Knott et al., 2008). Our method allows us to resolve function in large numbers of synaptic terminals in a single tissue block, where activity in each is quantifiable through a continuous three-dimensional readout of their functional vesicle pools. We process these image stacks using machine learning-based methods to automatically obtain segmentations of neurons, active zones, mitochondria and vesicles (classified as activated or not activated). We applied our method to investigate ultrastructure-function relationships in synapses activated by sensory input in primary visual cortex in awake head-fixed mice (**Fig. 1a**) (Marra et al., 2012). Synapses were labelled by intracranial infusion of FM1-43FX dye in layer 2/3 visual cortex circuitry (**Fig. S1**) via an access port embedded in a cranial window, as the animal simultaneously received a visual stimulus (either a drifting grating or darkness control). Since activated synapses take up dye into recycling vesicles, the internalized signal provides a direct report of presynaptic function (**Fig. 1b**). In tissue slices sectioned from the brain after fixation, the photostimulated fluorophores (**Fig. 1c,d**) drove diaminobenzidine photoconversion (**Fig. 1e**), yielding an electron-dense product that permitted investigation of functional vesicles in ultrastructure.

**Figure 1.**
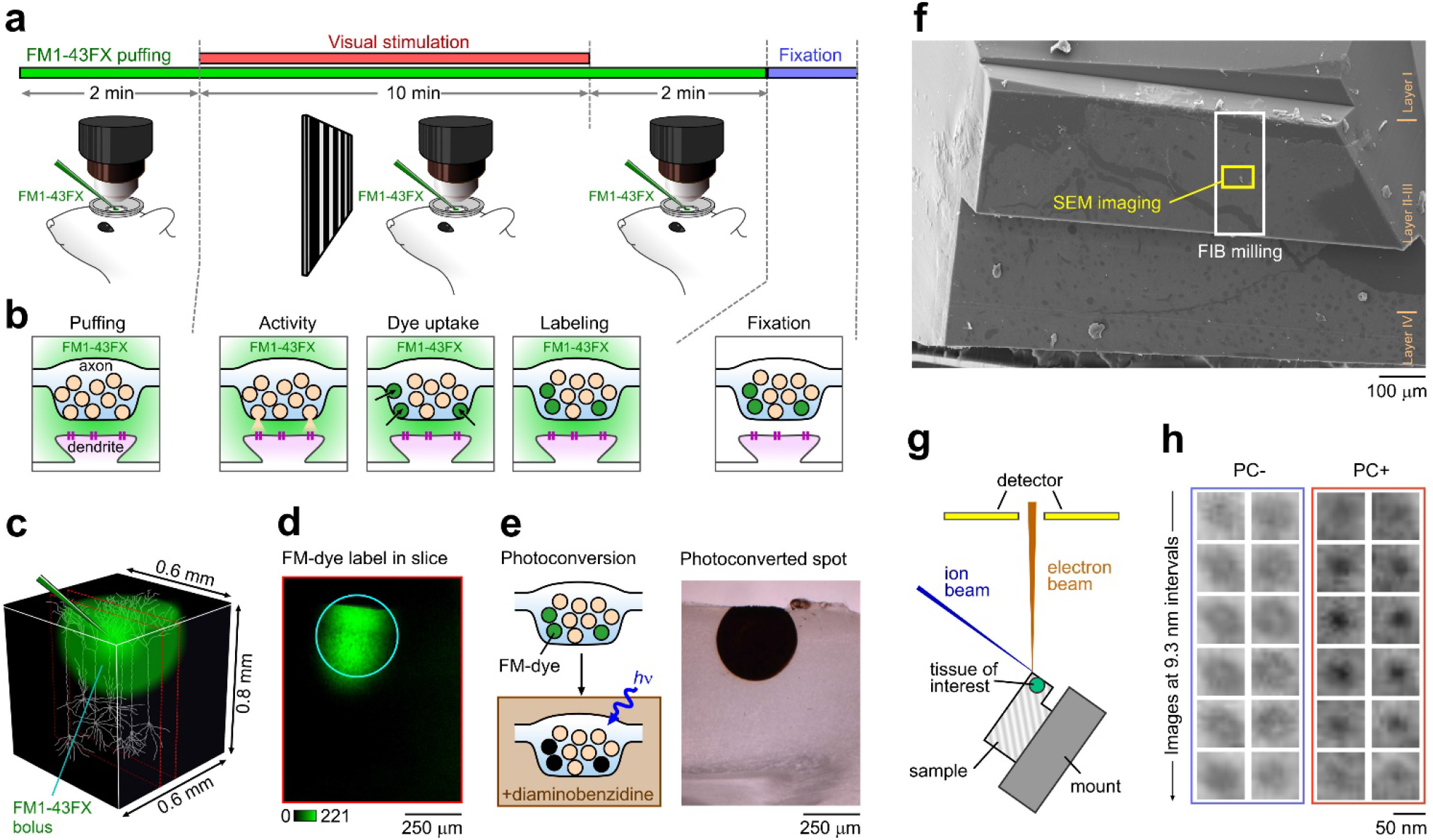
*In vivo* labelling of synaptic activity for ultrastructural investigation. **(a)** Schematic of experimental approach. FM1-43FX was injected via a patch pipette into Layer 2/3 of V1 in awake head-fixed mice running on a treadmill, while a drifting grating or dark control visual stimulus was delivered. **(b)** Cartoon illustrating mechanism of FM1-43FX dye-labelling of recycled vesicles. **(c)** Representation of FM-dye bolus in tissue (see also Fig. S1). Dashed red lines indicate a 100 μm slice taken from labelled region after fixation and imaged in **(d)** showing the FM-dye signal targeted for photoconversion. **(e)** (left) Schematic outlining the photoconversion process and (right) showing the tissue slice in (d) after photoconversion is complete. **(f)** Scanning electron microscope image of sample prepared for FIBSEM showing targeted region for imaging (yellow rectangle) and milling (white rectangle). Orange lines and text indicate cortical layers. **(g)** Schematic of FIBSEM setup. **(h)** FIBSEM images showing two distinct vesicle classes, non-photoconverted (PC-) and photoconverted (PC+) and their appearance through six consecutive sections (9.3 nm between sections).

Target regions were trimmed and prepared for FIBSEM (**Fig. 1f,g**) and high resolution 3D data acquired at 6000x magnification using a near-isotropic voxel size (6.2 x 6.2 x 9.3 nm). In this way, synaptic vesicles could be readily visualized in 5-6 consecutive sections (**Fig. 1h**). Strikingly, imaging in samples from the drifting grating stimulus condition revealed two distinct vesicle classes: photoconverted (PC+), characterized by an electron-dense lumen, and non-photoconverted (PC-, **Fig. 1h**), corresponding to recycled and non-recycled vesicles respectively (Marra et al., 2012). Spontaneous activity (darkness control) yielded PC+ vesicle counts of approximately one-tenth of those observed with visual stimulation. Control experiments confirmed that the labelling of vesicles was specifically due to the photoconversion of DAB in the presence of FM1-43FX (**Fig. S2**). PC+ vesicles persisted to depths > 100 μm from the photo-illuminated surface (**Fig. S3**), demonstrating that large tissue volumes are amenable to investigation with this method.

For visualization and analysis, we used convolutional neural networks to automatically segment neurons (Sheridan et al., 2021), active zones (Heinrich et al., 2018), mitochondria and vesicles. For the latter, we used a 3D U-Net (Ronneberger et al., 2015) to predict, for each voxel, whether it was part of a vesicle, and if so, whether it was photoconverted or not. Those predictions were then used to find the most likely positions of PC+ and PC-vesicles via a Hough transform with a spherical kernel (**Fig. 2** and Methods).

**Figure 2.**
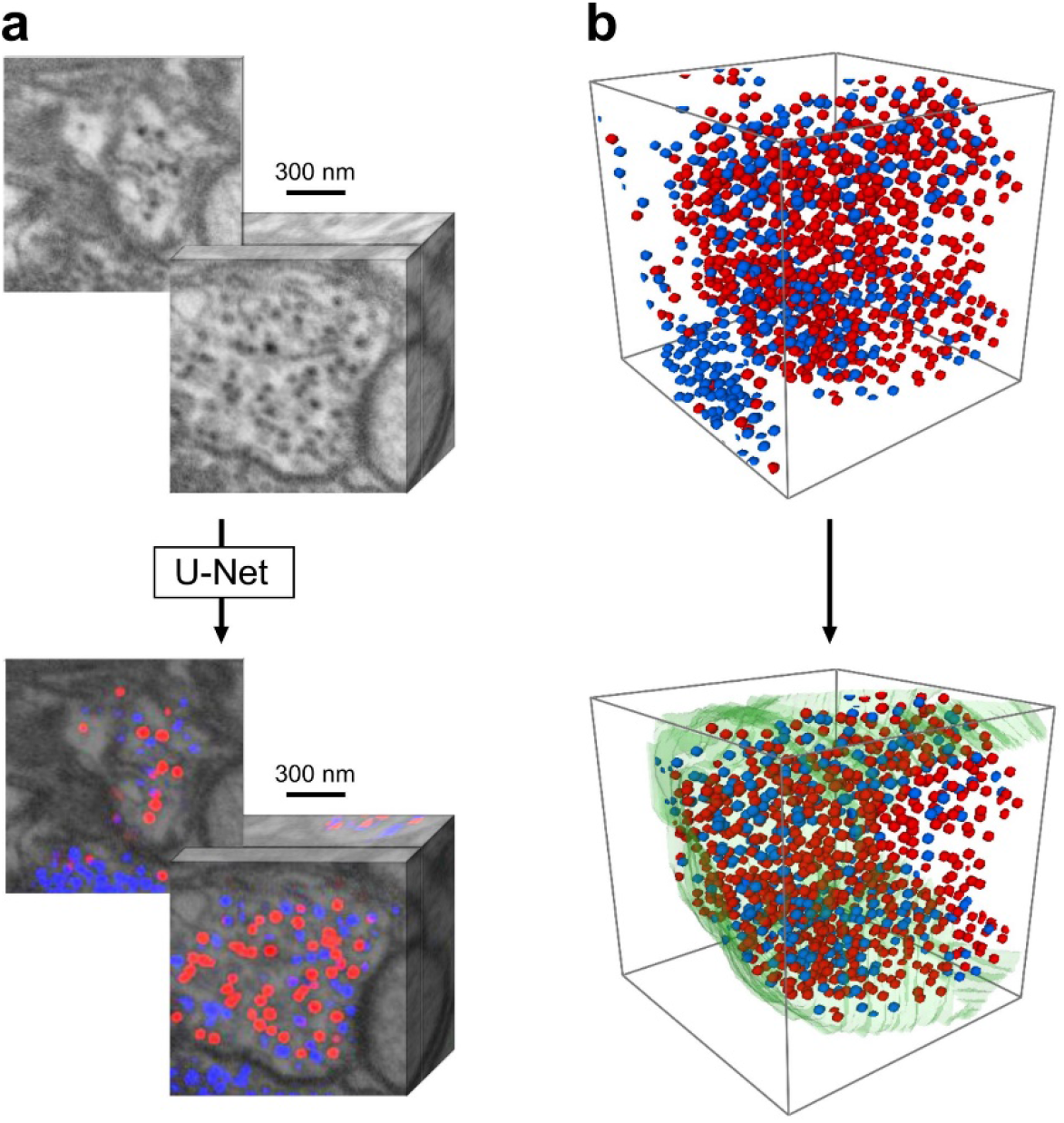
Illustration of the Hough transform vesicle detection on an example crop-out. **(a)** A 3D U-Net predicts, per voxel, the probability of being part of a PC+ (red) or PC- (blue) vesicle. **(b)** Convolution with a Hough kernel (sphere with diameter of 30 nm) followed by local non-max suppression yields a set of confidence-scored candidate detections, which are subsequently filtered to obtain non-overlapping, high confidence detections within a segmented bouton.

Outcomes of this segmentation are shown for a region of interest in cortical layer 2/3 from an animal presented with a drifting grating visual stimulus. In a single section from the raw FIBSEM stack (271 consecutive sections total, z-depth: 2.52 μm, **Fig. 3a**), PC- and PC+ vesicles are visible in synapses across the field of view with detail shown for two synaptic terminals in **Fig. 3b**.

**Figure 3.**
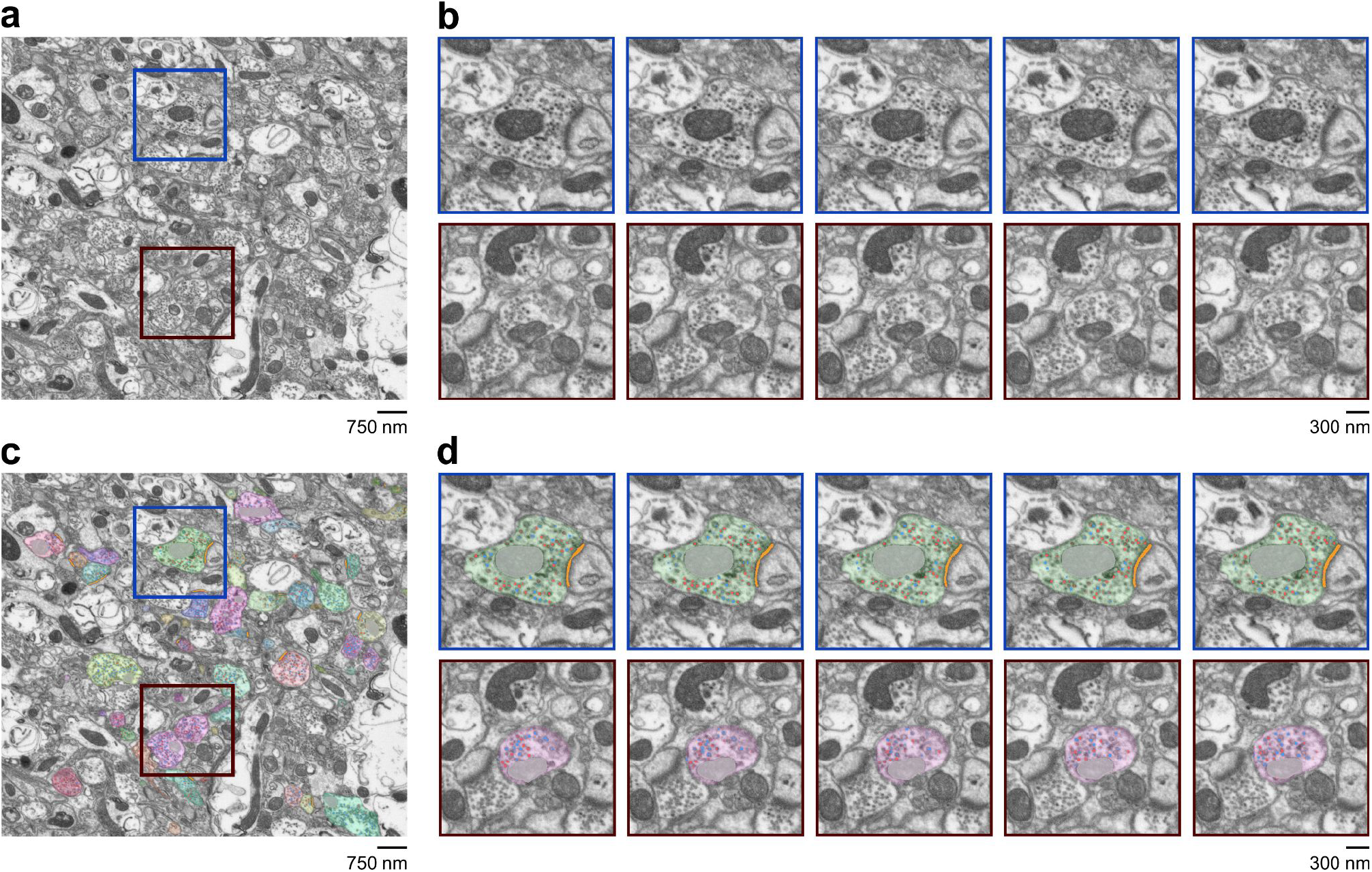
Ultrastructural readout of synaptic activity in a cortical tissue volume. **(a)** Sample image from FIBSEM stack (section 113/271) of layer 2/3 primary visual cortex from an animal that received a drifting grating visual stimulus to drive functional vesicle pool labelling. Coloured squares correspond to two synaptic boutons shown in detail in (b). **(b)** Top panel (synapse 1) corresponds to consecutive sections 106 - 110. The full bouton was present in sections 61 - 200 (1.29 μm in the z-plane). Bottom panel (synapse 2) corresponds to consecutive sections 83 - 87. The full bouton was present in sections 45 - 145 (0.93 μm in the z-plane). Both synapses are characterized by the presence of PC- and PC+ vesicles. **(c)** Same image as in (a) with coloured overlay showing outputs from detection, classification and segmentation steps. PC- and PC+ vesicles are classified as blue and red circles, respectively. Different axonal structures are assigned random colours. Mitochondria and active zones appear grey and orange, respectively. **(d)** Panels show the same images from (b) with coloured overlay as in (c).

The outputs of the detection, classification and segmentation steps were then added to the same images using coloured overlays, allowing characteristics of the ultrastructure and function to be directly assessed (**Fig. 3c,d**). Next, to view the segmented elements in the context of the whole tissue sample, we generated a complete 3D reconstruction and rendering of 109 axon volumes in the FIBSEM stack (**Fig. 4**) including the two boutons shown in Fig. 3b,d (**Fig. 4b,c**).

**Figure 4.**
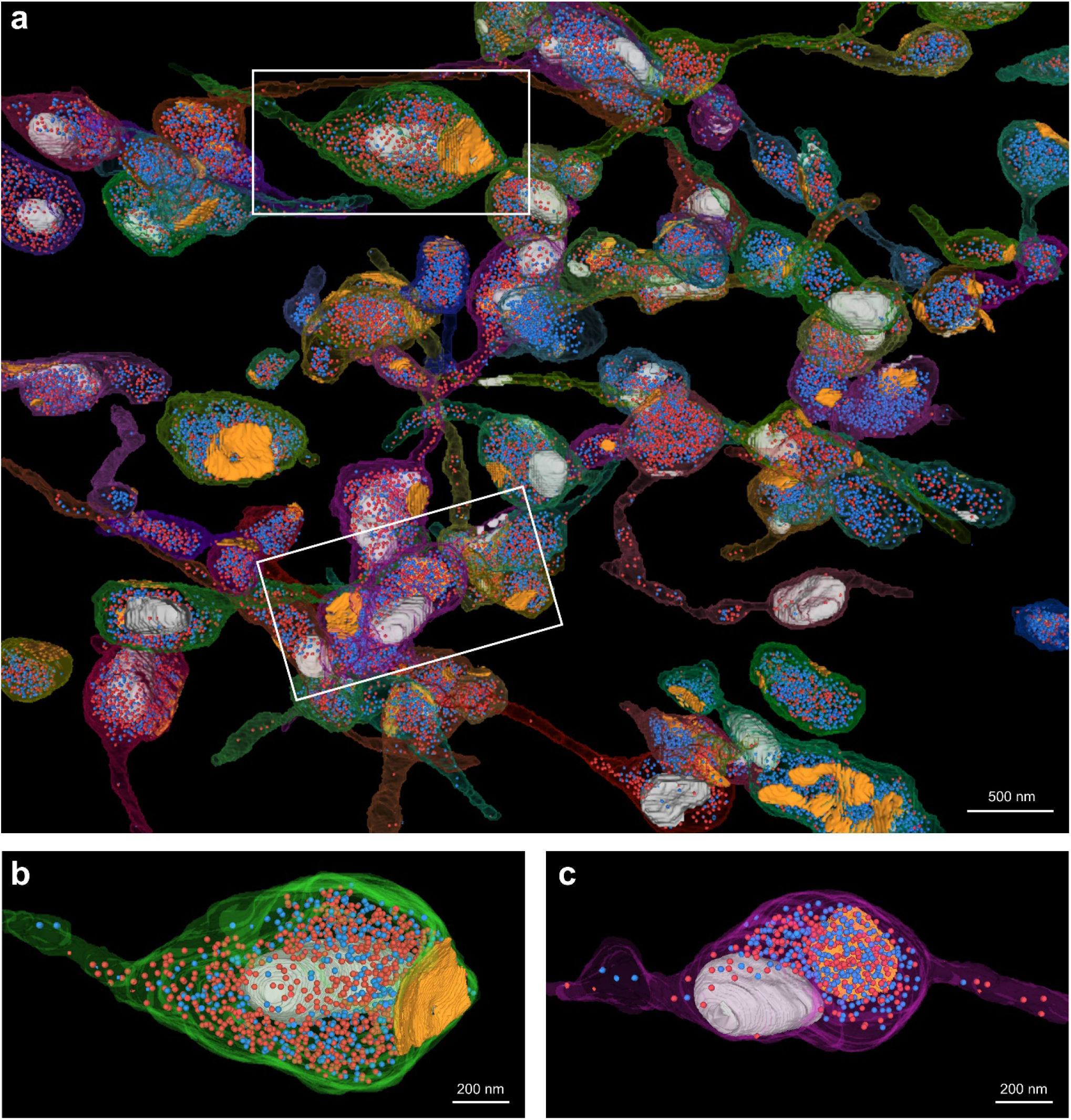
3D reconstructions of axon volumes in the FIBSEM stack. **(a)** 3D reconstruction from the FIBSEM stack shown in Fig. 3. 109 axons were reconstructed and displayed using Neuroglancer (opensource.google/projects/neuroglancer) with detail on a subset shown here. PC- and PC+ vesicles are classified as blue and red circles, respectively. Different axonal structures are assigned random colours. Mitochondria and active zones appear grey and orange, respectively. White rectangles correspond to synaptic boutons detailed in **(b and c)**.

For quantification, we counted both types of detected vesicles within the segmented axon volumes and assigned them to individual boutons based on their distance from its largest active zone. The distributions of the number of PC-vesicles (**Fig. 5a**), PC+ vesicles (**Fig. 5b**) and their sum (**Fig. 5c**) have a lognormal appearance, consistent with the idea that the brain employs lognormal distributions of firing rates and synaptic weights (Buzsaki & Mizuseki, 2014) for efficient signalling. In particular, a lognormal distribution of the number of PC+ vesicles is expected because the product of two random variables – spiking activity and release probability – both being lognormally distributed again follows a lognormal distribution. The number of PC+ vesicles scales with the total number of vesicles on average, yet the fraction of PC+ vesicles in the total vesicle pool varies greatly between individual synapses (**Fig. 5d**). For small boutons containing less than 50 vesicles, some of this variability may be contributed by shot noise in vesicular release, but also in boutons containing more than 50 vesicles a wide range of PC+ vesicle fractions was observed, consistent with the total number of vesicles being only a partial predictor of synaptic strength and activity (Branco et al., 2010). To distinguish variability in presynaptic spiking activity in different axons from variability in release probability of different boutons, we took advantage of the fact that some single axons in our FIBSEM stack contain multiple boutons (**Fig. 5e**), which therefore share the same number of presynaptic action potentials during the experiment. Differences in the numbers of PC+ vesicles in different boutons of the same axon, expressed as their ratio normalized to 1 (**Fig. 5f**), therefore allow us to assess their relative release probability. Strikingly, we find that the distribution of the ratios of PC+ numbers (**Fig. 5f**) is dominated by values of 0.4 or less, indicating that large differences in release probability in neighbouring boutons sharing the same presynaptic action potentials are common, and are consistent with a heavy-tailed distribution of presynaptic release probabilities. We illustrate this using a simple model in which release probabilities are lognormally distributed (**Fig. 5g,h**) and ratios of release probability are sampled in pairs and triples from this distribution (in the same proportion as the numbers of axons with two and three boutons analyzed in **Fig. 5f**). The resulting distribution of release probability ratios (**Fig. 5i**) peaks below 0.4, consistent with the experimental data, unlike release probability ratios obtained assuming normal or uniform distributions of release probability (data not shown).

**Figure 5.**
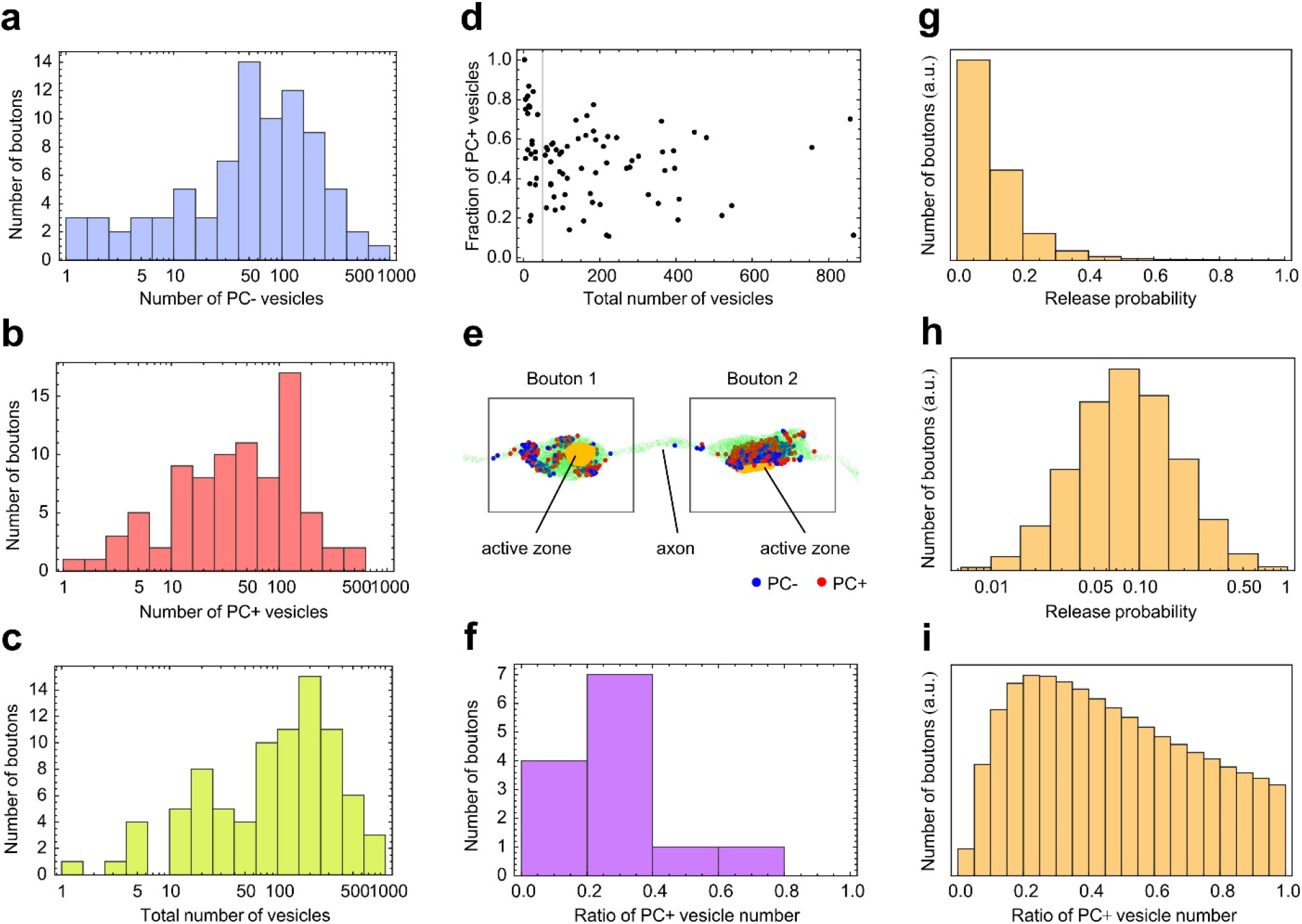
Quantifying presynaptic activity and release probability. **(a-c)** Histograms of the numbers of PC-, PC+ and total vesicles detected in n = 84 individual boutons. **(d)** Relationship of the fraction of PC+ vesicles and total number of vesicles. Above 50 total vesicles (vertical line), the PC+ fraction covers a broad but consistent range, independent of the total number of vesicles. **(e)** Volume rendering of an axon (green) with two boutons, each having one active zone (orange) and containing PC+ (red) and PC- (blue) vesicles. **(f)** Distributions of the ratios of the number of PC+ vesicles calculated for multiple boutons (n = 7 double boutons and n = 3 triple boutons) of the same axons, with the maximum number of PC+ vesicles normalized to 1 and not included in the histogram. In a model of lognormally distributed release probabilities **(g and h**; sampled a million times), release probability ratios sampled analogously to those in the data **(f)** follow a distribution that also peaks at low ratios **(i)**.

In summary, our approach provides an experimental strategy to quantify activity in synaptic boutons through large tissue volumes with full ultrastructural context. Given the near-isotropic imaging resolution that is achievable using FIBSEM, this offers a much more comprehensive 3D view of functional synaptic vesicle pools than conventional serial-section electron microscopy methods permit and can therefore yield detailed new insights into synaptic ultrastructure-function relationships. Our method measures the product of synaptic release probability and the number of action potentials evoked by sensory stimulation to give an estimate of total synaptic activity in each bouton. Furthermore, we demonstrate that by comparing different boutons belonging to the same axon, their relative release probabilities can also be assessed. A variant of this approach could tightly regulate imposed action potential activity, for example using all-optical stimulation methods (Packer et al., 2015), to estimate release probability directly at each terminal within a circuit of interest. Such a strategy would open up possibilities for elucidating the specific combinations of structural features of synapses and their environment that might be key predictors of synaptic strength.

## Methods

### Animals

All animal procedures were approved by the local Animal Welfare and Ethical Review Board at University College London and performed under license from the UK Home Office in accordance with the Animals (Scientific Procedures) Act 1986. Animals were maintained on a 12 h light/dark cycle with food/water ad libitum. We used C57BL/6 mice, 10 – 15 weeks old of both sexes without randomization. Mice were group housed before surgery and single-housed after surgery. In total, 8 mice were used to develop and optimize the *in vivo* labelling and EM approach, n = 5 were used to generate ultrastructural data (including validation controls) and from these, FIBSEM data was presented for n = 2 mice.

### Headplating and window installation

The installation of a cranial window was based on procedures described previously (Goldey et al., 2014). A minimum of 4 – 6 h before surgery, mice were injected i.m. with 4.8 μg g^-1^ dexamethasone (2 mg ml^-1^) to prevent cerebral oedema during surgery. Carprofen (Carprieve; 5 μg g^-1^) and buprenorphine (Vetergesic; 0.1 μg g^-1^) were administered s.c. peri-operatively and at 24 h intervals for 3 days for analgesia. Mice were anaesthetised using isoflurane (3 *—* 5 % for induction and 0.5 *—* 1.5 % for surgery with 0.5 *—* 1 liter min^-1^ O_2_) placed on a heating pad (37.5 ± 1 °C) and the head was stabilised in a stereotaxic frame (Kopf Instruments). The scalp was removed bilaterally from the midline to the temporalis muscles and a custom-made metal headplate with an 8 mm circular imaging window was fixed to the skull overlying the left primary visual cortex with dental cement (Super-Bond C&B, Sun-Medical). A 4 mm craniotomy centered 3.0 mm lateral to lambda, and 1.5 mm anterior to the lambdoid suture was performed to expose the V1 region of the visual cortex for window installation. A 4 mm coverslip (Menzel-Glaser) with a sealed (Kwik-Seal) laser cut window (0.5 x 1.5 mm) was press-fit in to the craniotomy, secured to the skull by a thin layer of cyanoacrylate (VetBond) and fixed in place by dental cement. The imaging well was coated with black dental cement for light blocking purposes and filled with Kwik-Cast to protect the window during recovery. Mice were allowed to recover for a minimum of 5 days before experimentation. Weight and body condition were monitored at regular intervals following surgery. One day before the experiment, a black light-proofing dish was secured on top of the imaging window under light anaesthesia. On the day of experiments, the Kwik-Cast was detached from the imaging well and the seal removed from the cranial window under brief isoflurane anaesthesia (< 20 min). Mice were allowed to recover for at least 20 min on the wheel before experimentation.

### *In vivo* visual stimulation and FM1-43FX bolus

20 μM FM1-43FX (Invitrogen) was freshly made with oxygenated aCSF on the day of the experiment from 10 mM stock and injected into L2/3 of primary visual cortex (120 *—* 170 μm) in awake head-fixed mice running on a treadmill. A long tempered small tip (2 *—* 3 μm in diameter) patch pipette (∼3 MΩ; borosilicate glass, WPI, GBF 150-86-10) was used to pressure inject the dye into the extracellular space at 4 *—* 6 psi for 14 minutes. After the initial 2 minutes of dye injection, a sinusoidal drifting grating with a spatial frequency of 0.04 cycles per degree and with a 2 cycles/s temporal frequency was delivered for 10 minutes (Smith et al., 2013). In control experiments, mice were kept in the dark instead of the grating stimulus for the same length of time (10 min). At the end of the period of visual stimulation, FM1-43FX injection continued for a further 2 mins. The FM1-43FX bolus location was imaged by 2-photon microscopy at the end of the protocol (bolus diameter: 350 *—* 450 μm, **Fig. S1**).

### Fixation and sectioning

At the end of the *in vivo* protocol, mice were terminally anaesthetized by intraperitoneal injection of ketamine (100 mg kg^-1^) and xylazine (15 mg kg^-1^) and transcardially perfused with 4% paraformaldehyde and 0.1% glutaraldehyde in 0.1 M phosphate buffer saline (PBS, pH 7.2) and post-fixed in the same fixative overnight at 4 °C. The brains were then sectioned on a vibratome (Leica VT1000S) at 100 µm and processed for photoconversion.

### FM1-43FX Photoconversion

The sections containing the FM1-43FX bolus were identified by mounting them onto slides and observing them under a fluorescence microscope. Any natural landmarks (e.g. blood vessels, notches in the tissue) were also documented in brightfield. These sections were then incubated in 100 mM glycine solution for 1 h, rinsed with 100 mM NH_4_Cl followed by a PBS wash (5 min). The sections were secured with a glass anchor in the bottom of a 35 mm petri dish and incubated in carbogen (5% CO_2_ and 95% oxygen) bubbled 3,3’-Diaminobenzidine (DAB; ACROS Organics; code: 328005000, CAS: 7411-49-6) solution (1 mg ml^-1^ in PBS) for 20 minutes in the dark. We replaced the incubation solution with fresh DAB solution and photoconverted the region with blue light (473 nm) from an LED source for 20 mins using a 40 x water immersion objective (ZEISS Achroplan, NA = 0.8) mounted on a custom made photoconversion microscope ((Dobson et al., 2019), 47 mW power) to allow effective FM dye photo-stimulation deep into the tissue (>100 μm, **Fig. S3**). This procedure was followed by rinses in PBS (3 x 1 min). The sections were then mounted on slides and imaged under an Olympus BX-51 microscope equipped with Neurolucida to document the location of the photoconversion spot and any of the previously identified natural landmarks to facilitate LM-EM correlation.

### Control experiments

Control experiments were performed to verify that the labelling of vesicles was only due to the photoconversion of DAB in the presence of FM1-43FX (**Fig. S2 a-c**, region 1), or when FM1-43FX was applied with DAB but not photoconverted (**Fig. S2 a-c**, region 2) or when no FM1-43FX was applied and no photoconversion was performed (**Fig. S2 a-c**, region 3). All TEM images are from samples prepared for FIBSEM (see below) and showed good ultrastructure preservation. Briefly, 70 nm ultramicrotome sections on Pioloform-coated copper slot grids were visualised with a FEI Tecnai T12 BioTwin transmission electron microscope at 120kV. Images were captured with an Olympus Morada CCD digital camera (magnification: 300x — 48000x), running iTEM software. TEM images from region 2 and 3 adjacent to region 1 which contains the photoconverted FM1-43FX bolus show a good ultrastructure and the absence of photoconverted vesicles in all synaptic terminals examined (**Fig. S2c**). Control experiments were also performed to determine the damage to the ultrastructure due to photoconversion as well as the extent of the photoconversion within the depth of the sections (**Fig. S3**). In both cases, TEM images from samples prepared for FIBSEM showed good ultrastructural preservation and the presence of photoconverted vesicles throughout the thickness of the section (**Fig. S3d**).

### FIBSEM: preparation

After photoconversion, sections were then processed for FIBSEM as in (https://ncmir.ucsd.edu/sbem-protocol) but in our case, replacing cacodylate buffer with 0.1 M phosphate-buffered saline (PBS, pH 7.4). The photoconverted sections were collected in PBS, postfixed in 1.5% (w/v) potassium ferrocyanide in 2% (v/v) osmium tetroxide in PBS (1 h, 4°C), followed by: 1% (w/v) thiocarbohydrazide in ddH_2_O (20 minutes, 60°C), 2% (v/v) osmium tetroxide in ddH_2_0 (30 minutes, RT), 1% (w/v) uranyl acetate (aqueous) (overnight, 4°C). Note each step was followed by ddH_2_O rinses. The sections were then dehydrated in an ascending scale of ethanol, followed by 100% acetone and infiltrated in 1:2, 1:1, 9:1 Durcupan epoxy resin:acetone for 2 h each. We then kept the sections in 100% Durcupan (10 g component A, 10 g component B, 0.3 g component D, 100 μl component C) at room temperature overnight. The following day, the sections were capsule embedded in freshly made 100% Durcupan resin and polymerised at 60 °C for 48 h. The region of interest in the photoconverted sections was trimmed for an initial semithin and then ultrathin sectioning with a Leica UC6 or UC7 ultramicrotome and visualized with a FEI Tecnai T12 BioTwin transmission electron microscope at 120 kV. To check photoconversion, images were captured with an Olympus Morada CCD camera (magnification: 300x – 48000x), running iTEM software. Blocks were then prepared for FIBSEM by reshaping the ROIs, placing them on stubs and silver painting (Agar Scientific) before sputtering them with gold.

### FIBSEM: Image acquisition

High resolution 3D data (6.2 x 6.2 x 9.3 nm voxels) at 6000x magnification were acquired using a focused ion beam scanning electron microscope (Zeiss NVision 40) controlled by custom software. This near-isotropic resolution was chosen to allow each synaptic vesicle to appear on 5-6 consecutive images. We used the 60 µA SEM aperture, SEM dwell time was 12.7 μs per pixel, and EHT was 1.50 kV. FIB milling was performed with I = 3 or 6.5 nA current at 30 kV as a compromise between milling speed and precision. All datasets were imaged with a single panel.

### Image registration and manual analysis: Fiji and Amira

All raw FIBSEM image stacks were registered in Fiji (http://fiji.sc/) using the Linear Stack Alignment with SIFT plugin. Amira (ThermoFisher) or MIB software (http://mib.helsinki.fi) was used for manual segmentation and rendering of axons with photoconverted vesicles within the same registered FIBSEM volumes also used for the automated detection of photoconverted vesicles (see below).

### Automatic Neuron and Mitochondria Segmentation

#### Neuron Segmentation

We used sparse ground truth consisting of 21 unique neurons (test = 11, control = 10) generated in Amira to train an initial 3D U-Net (same architecture as final U-Net, described below) to bootstrap further ground-truth generation. Once trained, we produced over-segmentations on several small (∼1 - 2 micron) cubes (test = 6, control = 6) and manually proofread them densely in Paintera (https://github.com/saalfeldlab/paintera) to segment neurons, glia, and plastic artifacts.

We then trained a final 3D U-Net based on a previously published architecture (Funke et al., 2019). The network learned a 10D embedding describing local object shape to aid in the prediction of a 3D affinity graph (Sheridan et al., 2021). We taught the network to learn to predict zero affinities in areas of the ground-truth which were labelled as glia and artifacts to prevent false merges between neurons.

Training was implemented using the gunpowder (https://funkey.science/gunpowder) library to augment the data with random flips (all dimensions), transposes (limited to the x and y dimensions), rotations (up to 90°), intensity scale (between 0.9 and 1.1) and shift (between - 0.1 and 0.1).

Boundaries were generated from the affinities for the test and control volumes. Supervoxels were created using a seeded watershed approach, and edges were iteratively merged using hierarchical region agglomeration with a 75th quantile merge function up to a certain threshold (Funke et al., 2019). We used a low agglomeration threshold (test = 0.1, control = 0.01) to obtain a segmentation in which false splits were more likely than false merges. Targeted proofreading was then performed on a subset of neurons (test = 109, control: 16) which were used for analysis.

#### Mitochondria Segmentation

Segmentation of mitochondria followed the same procedure as for the neurons. To this end, we manually annotated mitochondria in the test dataset (n = 21) and treated anything else as background.

### Automatic Detection of Active Zones

We used 8 volumes (test = 4, control = 4) with an average size of 5.84 μm^3^ and 7 active zones for training. We trained a 3D U-Net (12 initial feature maps, three max-pooling downsampling steps with size 1 x 2 x 2 in the first layer and 2 x 2 x 2 in the next two layers, and a feature map increase with factor 5 after each downsampling step) to predict the signed distance to the closest cleft (positive outside, negative inside), loosely based on a previous method (Heinrich et al., 2018), i.e., the sigmoid of the Euclidean distance. The predictions were then thresholded at 0.5 (corresponding to a signed Euclidean distance of zero or less) to obtain a segmentation. Augmentations were consistent with the neuron and mitochondria training pipelines.

### Automatic Vesicle Detection

Similarly, we trained a convolutional neural network to detect vesicles (PC+ and PC-). For that, we densely labelled all vesicles as PC+, PC-, and “uncertain” in a 2.79 x 4.31 x 1.86 μm^3^ region of interest using Ilastik (https://www.ilastik.org) or Amira for training, and a second region of size 3.72 x 2.01 x 0.28 μm^3^ for validation. We trained an initial 3D U-Net (48 initial feature maps, two max-pooling downsampling steps with size 2 x 2 x 2, feature map increase with factor 5 after each downsampling step) to classify each voxel into PC+, PC-, or background using a cross-entropy loss. To account for the class imbalance between background and vesicles, we scaled the loss contribution of background voxels with a factor of 0.01. We then used the initial network to label all “uncertain” vesicles in the training dataset as PC+ or PC-in a consistent manner and repeated training to obtain the final vesicle prediction network (same architecture as initial network).

Training was implemented with gunpowder using the same augmentations as the neuron segmentation pipeline. The trained network predicts, for each voxel, the probability for each of the three classes. In a post-processing step, those predictions were converted into vesicle detections by performing a Hough transform. To this end we convolved the sum of predicted probabilities for PC+ and PC-vesicles with an isotropic ball with diameter of 30 nm, followed by local non-max suppression with a minimum distance of two voxels between maxima. The maxima were considered as candidate detections, with their values as detection scores. We then greedily accepted vesicle candidates as true detections in descending detection score order until a given threshold (≥ 20, empirically determined on the validation dataset), ensuring that no spatially overlapping candidates were accepted at the same time. We further labelled each accepted vesicle as PC+ or PC-. For that, we considered the sum of predicted probabilities for PC+ and PC-of all voxels that belong to the detection and assigned the label based on the larger sum.

### Vesicle pool quantification

Analysis of vesicle pools filtered by the segmented axon volumes was performed in Mathematica. Vesicles counted within up to 0.8 µm from the largest active zone of a bouton were assigned to that bouton. For single-bouton analyses, data from axons containing a single bouton, and data from one randomly chosen bouton from each multi-bouton axon were pooled.

## Acknowledgements

We thank L. Goetz (WIBR, UCL) and S. Rey (Sussex, UK) for help with preliminary experiments, Zihui Zhang (WIBR, UCL) for assistance in setting up the photoconversion microscope, Julie Anh-Ton for help with tracing work, Jemima Burden and the MRC LMCB, EM Facility, UCL (MC U112266B) for support with sample preparation for TEM. This work was funded by the Wellcome Trust (201225/Z/16/Z to MH, 208348/Z/17/Z to KS, 108201/Z/15/Z to VM), the European Research Council (ERC-695709 to MH), UKRI (BBSRC, BB/K019015/1 and BB/S00310X/1 to KS) and the Gatsby Foundation (GAT2919 to MH).

## Author contributions

KS and MH conceived the work and acquired funding. AS, MF, CR and AR carried out experiments, advised by VM. JF, A.Sh and AR developed software. AR, AS, A.Sh, CR and JF analyzed the data. AS, CR and A.Sh carried out data segmentation and validation. AR, A.Sh, JF and KS produced visualizations. AR, CR, JF, KS and MH supervised the work. AR, JF, KS and MH drafted the manuscript. All authors reviewed and edited the manuscript.

