## Supplementary Figures for "Ultrastructural readout of *in vivo* synaptic activity for functional connectomics"

**Three supplementary figures:**

Figure S1

Figure S2

Figure S3

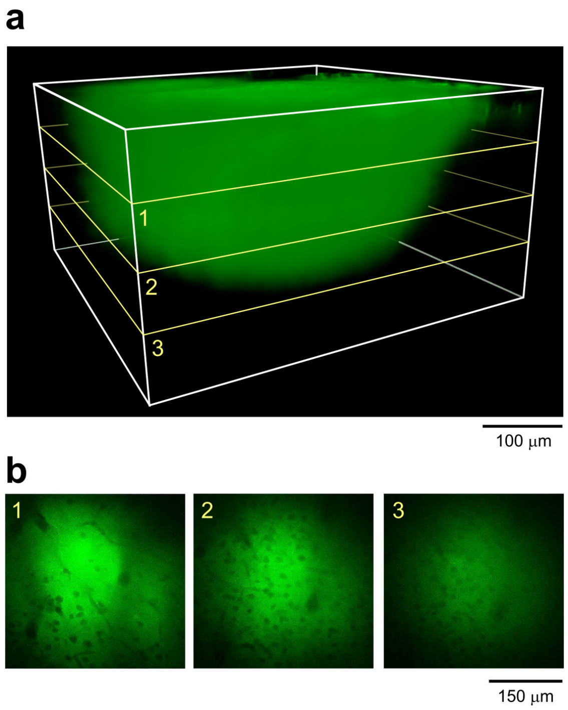


**Fig. S1. 3D imaging of FM1-43FX bolus used for synaptic labelling. (a)** 3D reconstruction of FM-dye bolus created from a 2-photon image stack collected at the end of the labelling protocol. **(b)** Slices 1, 2 and 3 from image stack in (a) showing homogeneous fluorescence intensity.

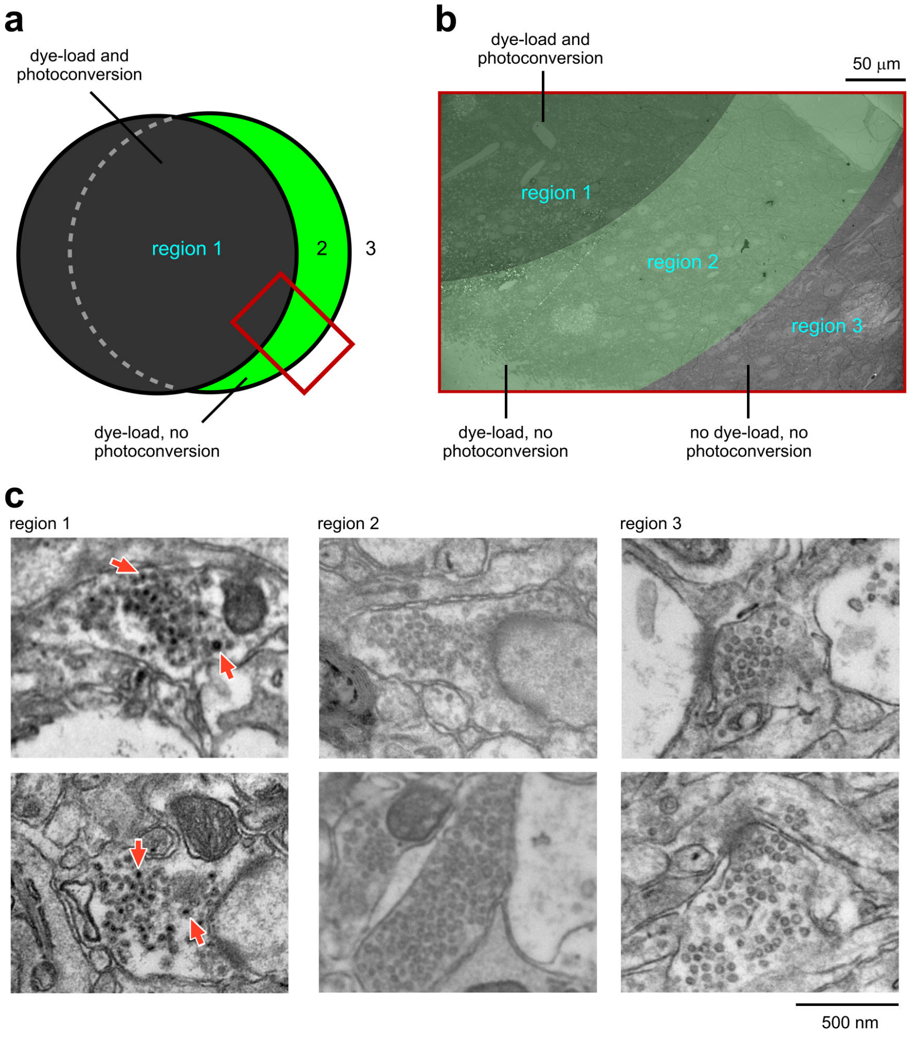


**Fig. S2. Vesicle labelling depends on photoconversion of diaminobenzidine in the presence of FM1-43FX. (a)** Schematic illustrating control experiment in which the photoconversion spot is displaced laterally from the region of FM-dye fluorescence, generating three regions: (1) dye-loading with photoconversion, (2) dye-loading, no photoconversion, (3) no dye-loading, no photoconversion. **(b)** Electron micrograph showing the three regions (red rectangle in a). **(c)** Example electron micrographs of each synapse in each region. Region 1 is characterized by PC+ vesicles (red arrows) while regions 2 and 3 are not.

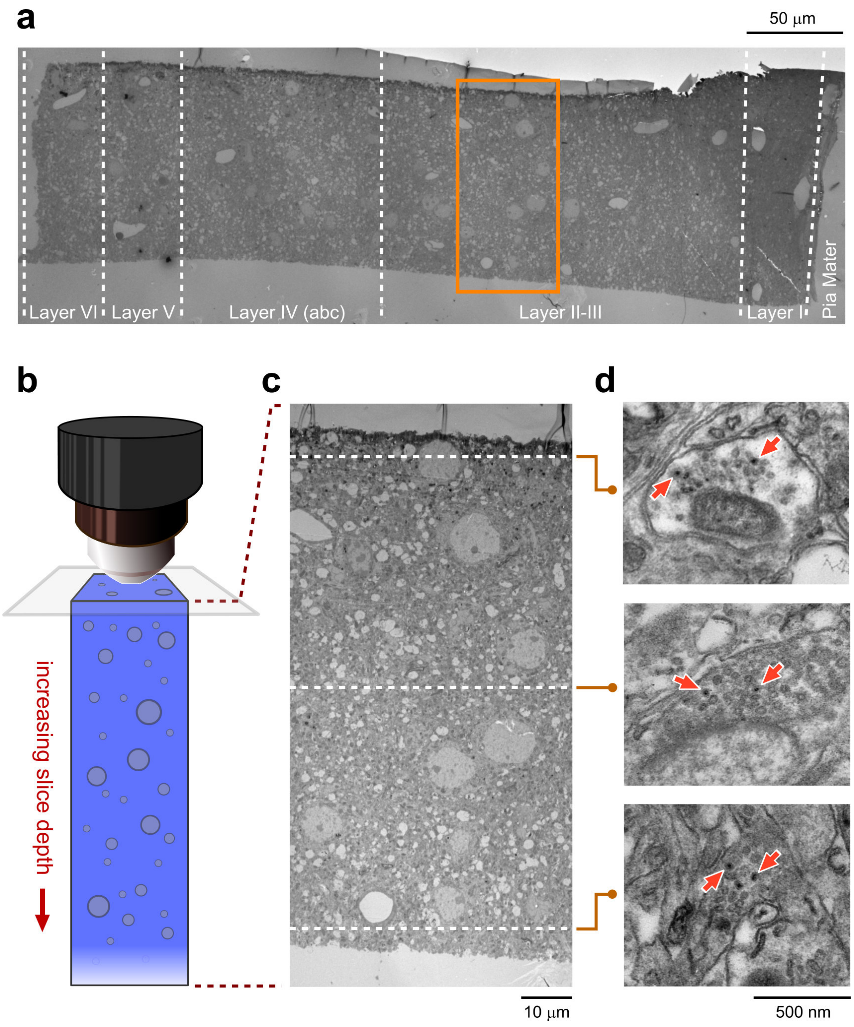


**Fig. S3. Reliable FM-dye photoconversion persists deep (> 100 µm) into the tissue slice.** **(a)** Low magnification electron micrograph shows cortical layers and region targeted (orange rectangle). **(b)** Schematic illustrates the photoconversion setup where 473 nm light from an LED source is used to illuminate the sample. **(c)** Detailed view of region targeted for photoconversion. **(d)** Consistent FM-dye labelling was seen at all depths away from the photo-illuminated surface.
